# A Comparison of Deep Neural Networks for Seizure Detection in EEG Signals

**DOI:** 10.1101/702654

**Authors:** Poomipat Boonyakitanont, Apiwat Lek-uthai, Krisnachai Chomtho, Jitkomut Songsiri

## Abstract

This paper aims to apply machine learning techniques to an automated epileptic seizure detection using EEG signals to help neurologists in a time-consuming diagnostic process. We employ two approaches based on convolution neural networks (CNNs) and artificial neural networks (ANNs) to provide a probability of seizure occurrence in a windowed EEG recording of 18 channels. In order to extract relevant features based on time, frequency, and time-frequency domains for these networks, we consider an improvement of the Bayesian error rate from a baseline. Features of which the improvement rates are higher than the significant level are considered. These dominant features extracted from all EEG channels are concatenated as the input for ANN with 7 hidden layers, while the input of CNN is taken as raw multi-channel EEG signals. Using multi-concept of deep CNN in image processing, we exploit 2D-filter decomposition to handle the signal in spatial and temporal domains. Our experiments based on CHB-MIT Scalp EEG Database showed that both ANN and CNN were able to perform with the overall accuracy of up to 99.07% and F1-score of up to 77.04%. ANN with dominant features is more capable of detecting seizure events than CNN whereas CNN requiring no feature extraction is slightly better than ANN in classification accuracy.

## I. INTRODUCTION

An epileptic seizure is defined by the International League Against Epilepsy (ILAE) as a transitory occurrence of signs and/or symptoms due to abnormal excessive or synchronous neuronal activity in the brain [1]. It was reported that 65 million people of all ages have the epilepsy [2]. Owing to the impacts of epileptic seizures, which can lead to neuronal injuries, patients with recurrent or prolonged seizures should be reviewed by neurologists for a prompt diagnosis and treatment [3], [4]. For those with refractory status epilepticus unresponsive to medication, neurologists usually monitor the patients with continuous video-EEG monitoring [5], [6]. This is a combination of EEG and video, recorded simultaneously to observe brain activities in correlation with a clinical change. Nevertheless, this task is still a time-consuming process for the neurologists to review the continuous EEG. Therefore, an automated epileptic seizure detection using EEG signals is developed to facilitate the interpretation of long-term monitoring [7].

Ictal EEG pattern is a sequence of spike, sharp wave, and slow wave when seizures occur [8]. By the morphology of these three patterns, there are continuous changes in amplitude, frequency, and rhythms relative to the background [9]. When a seizure originates at some point and rapidly engaging the whole networks, causing EEG changes apparently appear on the whole brain, it is called a generalized-onset seizure. On the other hand, a seizure is focal-onset when originating within networks limited to one hemisphere, making the EEG changes restricted in a particular brain region [10], [11].

Many researchers have applied machine learning techniques in the automated seizure detection [12]. Local Binary Pattern (LBP) was used to reflect the differences of patterns from normal and ictal EEG signals using k-nearest neighbor (k-NN) [13]. Similarly, Local Neighbor Descriptive Pattern (LNDP), and Local Gradient Pattern (LGP) were developed to overcome a disadvantage of LBP which is locally invariant [14]. When combined with k-NN, support vector machine (SVM), decision tree (DT), and ANN, LNDP and LGP are able to classify each sample of EEG time series into seizure and normal group successfully. Analysing EEG signals in frequency domain is also common. The first four minimum and maximum amplitudes, and their corresponding frequencies were extracted from the power spectral density of EEG signal to measure frequency properties in seizure [15]. All extracted features were then applied to Gaussian mixture model, ANN, and SVM for classification. Weighted permutation entropy (WPE) was computed from coefficients of discrete wavelet transform [16]. WPE from each decomposition level was concatenated into a feature vector and the vector was fed into three classifiers, namely, linear SVM, radial basis function kernel SVM (RBF SVM) and ANN. Some study attempted to select features based on relevance and redundancy analysis in seizure detection to reduce computational complexity [17]. Although these feature-based methods achieved typically promising results, they still require background knowledge about seizure characteristics in the feature extraction process.

On the other hand, deep learning approaches can be exploited to seemingly characterize and recognize a nature of EEG signals. Various purposes of applying CNNs include classifying spectrogram of a small EEG epoch [18] using a modified stacked sparse denoising autoencoder (mSSDA), classifying seizure occurrences from a single-channel intracranial EEG signal [19], generating suitable features for interictal epileptiform discharges (IED) detection [20], or detecting interictal epileptiform spikes from scalp EEG recordings [21].

However, to our knowledge, there has been no research focusing on a deep CNN in detecting seizure using *raw scalp* EEG signals from *all channels*. It is more likely that common epileptic cases contain generalized seizures so a detection process using signals from all channels can become a common practice and require no prior knowledge about EEG characteristics. Hence, this paper aims to explore a capability of a deep CNN model in seizure detection using raw scalp EEG signals and compare its performance with a deep ANN model using multiple selected features as inputs. We also present a feature selection based on the Bayesian classifier [22] to choose dominant features independently. The performances of deep CNN and ANN models will be evaluated based on accuracy and F1-score of classification tasks and tested on the public CHB-MIT database.

This paper is organized as follows. Firstly, we describe the significance of features based on the Bayesian theorem and feature relevance in Section II. The description of database, subject demographic and the process of signal segmentation are explained in Section III. This section also presents the proposed scheme of applying CNN model and describe ANN structures. Numerical results and discussions are shown in Section V.

## II. FEATURES FOR SEIZURE DETECTION

In seizure detection, there are many features, e.g., statistical parameter, energy, and entropies [23], commonly used with machine learning techniques. However, using too many insignificant and relevant features requires more complex models and leads to high computational complexity and overfitting problem. For this reason, we apply a Bayesian-based method to determine significance of each feature and select only relevant features as inputs of ANN. Feature candidates calculated from time, frequency, and time-frequency domains are shown in Table I where features from time, frequency, and time-frequency domains are computed from a raw EEG signal, Fourier transform of the EEG signal, and coefficients of discrete wavelet transform (DWT), respectively. In DWT, EEG signals are decomposed into 5 levels using Daubechies 4 wavelet. Each feature is calculated on each channel simultaneously and the feature is averaged over all the channels from the left and right sides of the brain.

**TABLE I:** List of feature candidates.

| Domain | Feature |
| --- | --- |
| Time | Mean, Variance, Coefficient of variation, Skewness, Kurtosis, Max, Min, Energy, Nonlinear energy, Line length, Shannon entropy, Approximate entropy, Sample entropy, Number of zero crossing, Number of local extrema, Mobility, Complexity |
| Frequency | Intensity weighted mean frequency, Intensity weighted bandwidth [24], Spectral entropy, Peak frequency, Peak amplitude |
| Time-frequency | Mean, Absolute mean, Variance, Skewness, Kurtosis, Max, Min, Energy, Line length |

From literature [17], [24], these candidates are selected according to abilities to capture changes of amplitude, frequency, and rhythms. Figure 1 shows examples of feature that change corresponding to the changes in EEG signals. The dash and dashdotted lines indicate seizure onset and offset from an annotation, respectively. We can see that variance and nonlinear energy are typically high corresponding to the high amplitude in the EEG signal during the seizure activity. On the other hand, Shannon entropy in seizure duration is slightly different from the background. These suggest us that each feature responds to EEG signal differently; thus, combining many features can be more beneficial than applying an individual feature.

**Fig. 1:**
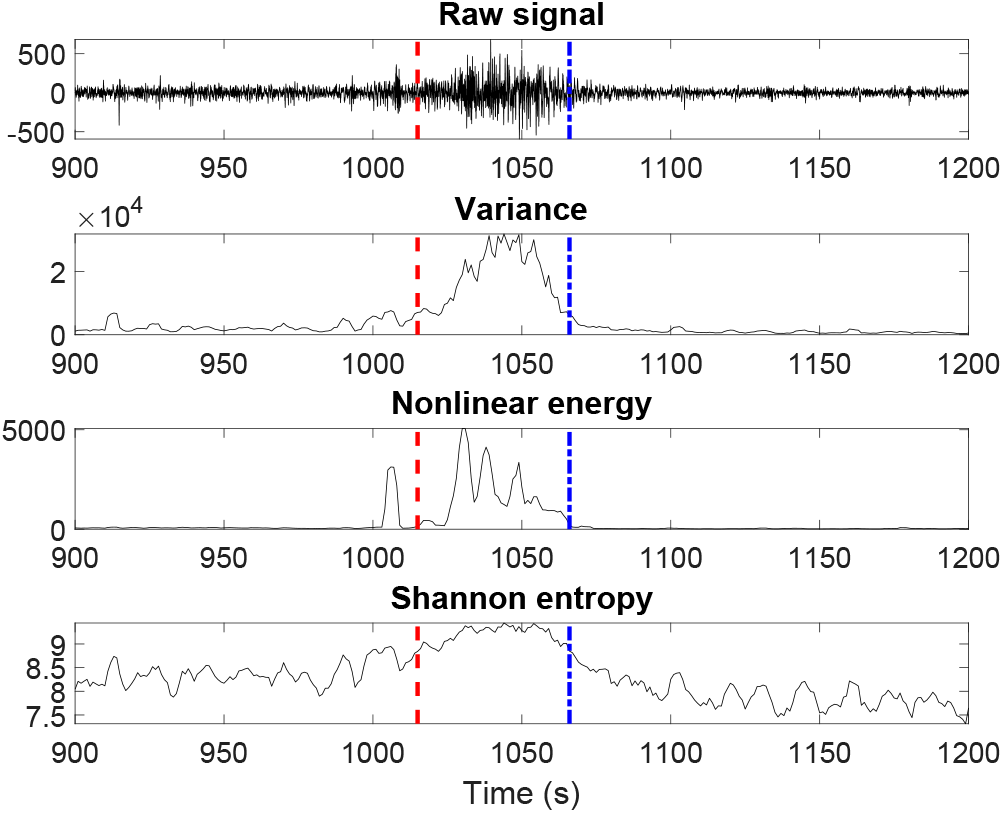
Feature examples that respond to changes of EEG signals. Each feature is calculated from 4-second EEG epochs and the sliding window is 1 second. This displayed signal was collected from the file *chb01-16* on the channel *FP1-F7*. Dash line indicates seizure onset and dashdotted line shows seizure offset.

The posterior probability density functions of the averaged features on the left and right sides are separately estimated using Gaussian kernel non-parametric density estimation [22]. Consequently, we apply the Bayesian classifier to each posterior probability function to obtain a classification error. However, using the Bayesian error rate could be misleading in the context of unbalanced data, especially in clinical diagnosis since data contain only a small fraction of seizure samples. For instance, the ratio of a number of abnormal samples to the normal sample size is 0.0001, the Bayesian error rate is then typically at most 0.0001, referring to the accuracy of at least 99.99%. This too optimistic result can be simply achieved by detecting every sample as normal. Hence, a score indicating an improvement of the Bayesian error from a baseline is used to determine the significance of each features, defined by

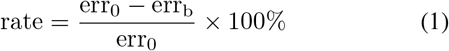

where err_b_ is the error from the Bayesian classifier and err_0_ is the error from the baseline method that classifies every segment as normal.

## III. SEIZURE DETECTION SCHEME

The proposed classification process for a deep ANN-based model is shown in Figure 2. The ANN model contains 7 hidden layers and each layer has 8 neurons. The chosen features from the Bayesian method are extracted from each channel separately. Consequently, a feature vector as the input to the ANN model is constructed by concatenating all features from all channels.

**Fig. 2:**
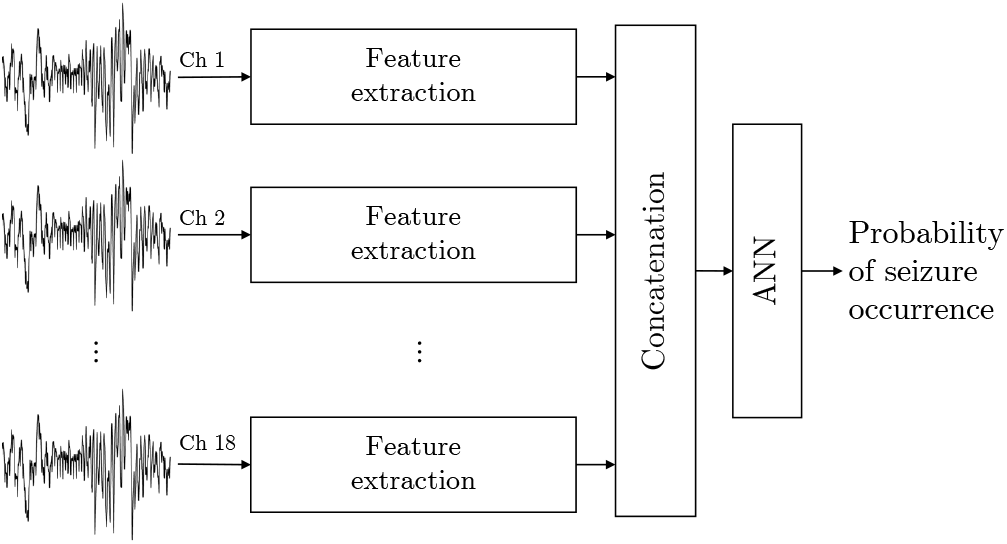
Classification process for ANN model. Chosen features are independently extracted from 18 channels and stacked into a feature vector. Then the feature vector is applied to ANN to determine a seizure activity.

CNN architecture in this study consists of convolutional layers, activation layers, normalization layers [25], pooling layers, dropout layers [26], and fully-connected layers. We propose a detection scheme using raw EEG signals from all channels as input as illustrated in Figure 3 and the proposed CNN model is depicted in the block where Conv(*h, w, f*) stands for a convolutional layer of size (*h, w*) with f filters, BN is a batch normalization layer, ReLU is a rectified linear unit as an activation layer, Max(*h, w*) is a max-pooling layer of size (*h, w*), Dropout(*α*) is a dropout layer with a fraction of *α* to disconnect the input units, and FC(*a, b*) stands for a fully-connected layer that receives *a* inputs, excluding *a* bias term, and produces *b* outputs. In order to capture ictal patterns in EEG signals with a convolutional layer, a filter width corresponding to an EEG signal in one channel should be higher than the filter height, meaning that *w* > *h*. The model is adjusted by reducing a number of filters and adding convolutional layers to prevent an overfitting problem while maintaining a high classification accuracy since the deeper model with less parameter is approximately better than the shallow model [27]. In addition to decreasing model complexity and parameters, the convolutional layer is decomposed into (*w*, 1) and (1, *h*) filters [28]. Moreover, a 1-by-1 convolutional layer [29] is added in order to reduce computational complexity.

**Fig. 3:**
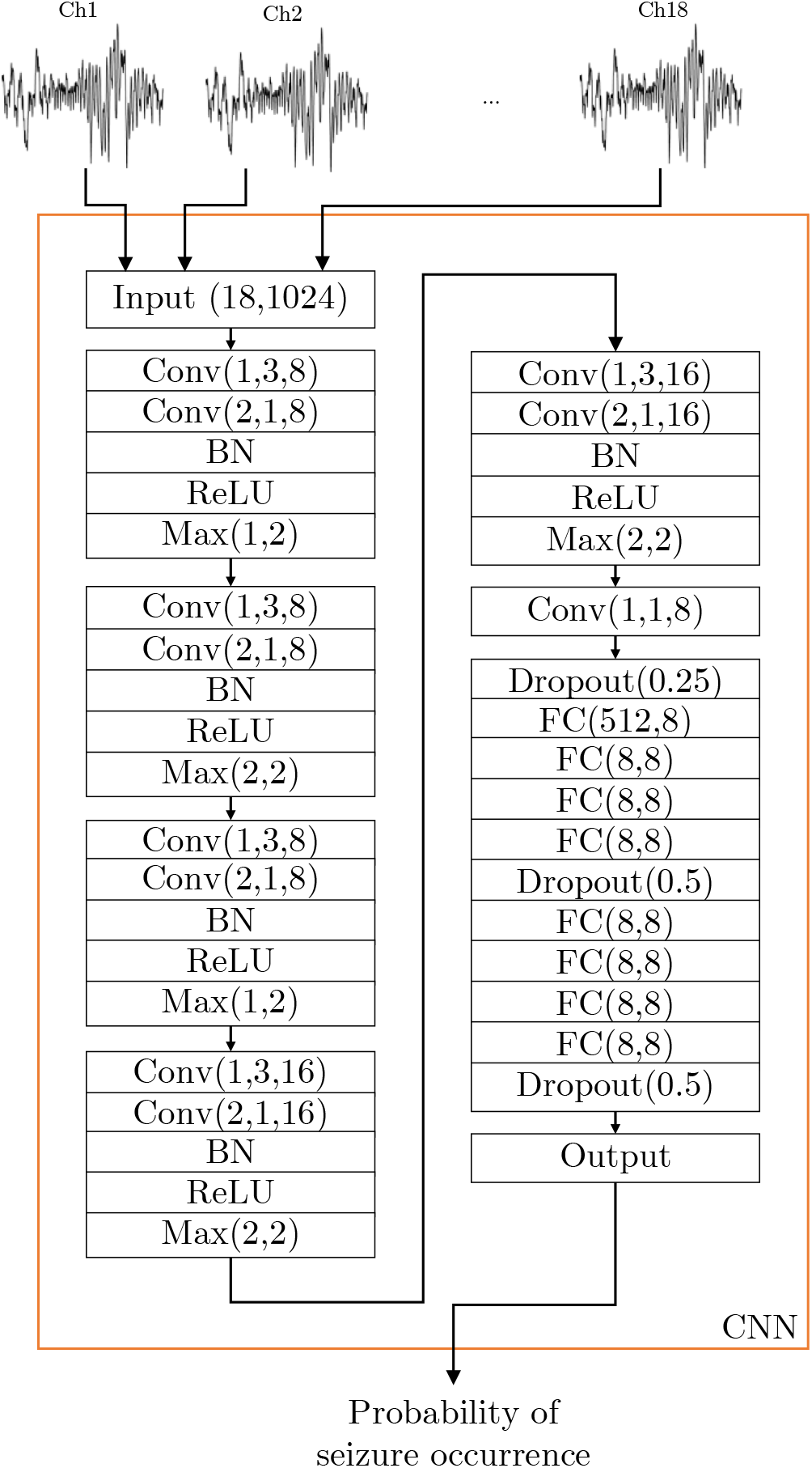
Classification process for the CNN model. Raw EEG signals from every channels are simultaneously fed to a convolutional neural network to produce the probability of seizure appearance.

## IV. DATA DESCRIPTION

This study uses the public CHB-MIT Scalp EEG Database [30] that consists of 24 EEG recordings from 23 patients: 5 males aged 3-22 years, 17 females aged 1.5-19 years, and one anonymous subject. All EEG data were obtained at Children’s Hospital Boston and fully deidentified and privacy-protected. All signals were measured with sampling frequency of 256 Hz at 16-bit resolution and stored in EDF file [31]. The international 10-20 system is used to locate electrode positions. This database is available online at PhysioNet (https://physionet.org/physiobank/database/chbmit/).

In our experiment, 24 records listed Table II were randomly selected based on the conditions that every record must have at least one seizure and was collected with a bipolar montage. According to the bipolar montage, the channels are FP1-F7, F7-T7, T7-P7, P7-O1, FP1-F3, F3-T3, T3-P3, P3-O1, FP2-F4, F4-C4, C4-P4, P4-O2, FP2-F8, F8-T8, T8-P8, P8-O2, FZ-CZ, CZ-PZ. So the total seizure duration from all chosen records is 2401 seconds and the total record duration is 182957 seconds; thus there exists 1.31% seizure period. We use each record separately to train and test the models. In every channel, each record is then segmented into samples with 4-second width, 1024 sample points per segment, and the moving-window is set to 1 second, with 3-second overlapping with the previous segment. As a result, the input of the CNN model has the size of 18 × 1024.

**TABLE II:** List of records.

| Records |  |  |  |  |
| --- | --- | --- | --- | --- |
| chb01_04 | chb01_16 | chb02_16+ | chb02_19 | chb03_03 |
| chb03_35 | chb04_08 | chb04_28 | chb05_06 | chb05_13 |
| chb06_01 | chb06_04 | chb07_13 | chb07_19 | chb08_02 |
| chb08_05 | chb09_06 | chb09_08 | chb10_38 | chb10_89 |
| chb23_06 | chb23_09 | chb24_04 | chb24_11 |  |

## V. RESULTS AND DISCUSSION

Features in Table I are computed from each EEG segment of each record and features from all conditionally random records are used to estimate posterior probability distributions and calculate improvement rates. Any feature with improvement rate greater than 5% is considered to be significant in seizure detection. As a result, time-domain features with high improvement rates shown in Table III were variance, energy, nonlinear energy, and Shannon entropy, whereas the other time- and frequency-domain features obtained apparently small improvement rates, less than 1%.

**TABLE III:** Bayesian error rate and improvement of features with high improvement rates on **left** and **right** sides of the brain.

| Feature | Left side |  | Right side |  |
| --- | --- | --- | --- | --- |
|  | err<sub>b</sub> | rate | err<sub>b</sub> | rate |
| Variance | 0.0156 | **7.76** | 0.0160 | **5.07** |
| Energy | 0.0156 | **7.76** | 0.0160 | **5.06** |
| Nonlinear energy | 0.0153 | **9.49** | 0.0154 | **8.69** |
| Shannon entropy | 0.0168 | 0.31 | 0.0159 | **5.85** |

Moreover, as shown in Figure 4, features calculated from wavelet coefficients which obtained high improvement rates were variance and energy from all decomposition levels, and line length from approximation coefficients of level 5. However, since variance and energy have similar contribution by their mathematical expression, causing redundancy, we neglected the energy from both time and time-frequency domains as a feature. So, instead of applying all 76 features, only 10 features are used and calculated from each channel independently. The dominant features used in further experiment with a deep ANN are listed as follows:

- **Time domain**: Variance, nonlinear energy, Shannon entropy.
- **Time-frequency domain**: Variance from all decomposition levels, line length from approximation level 5.

**Fig. 4:**
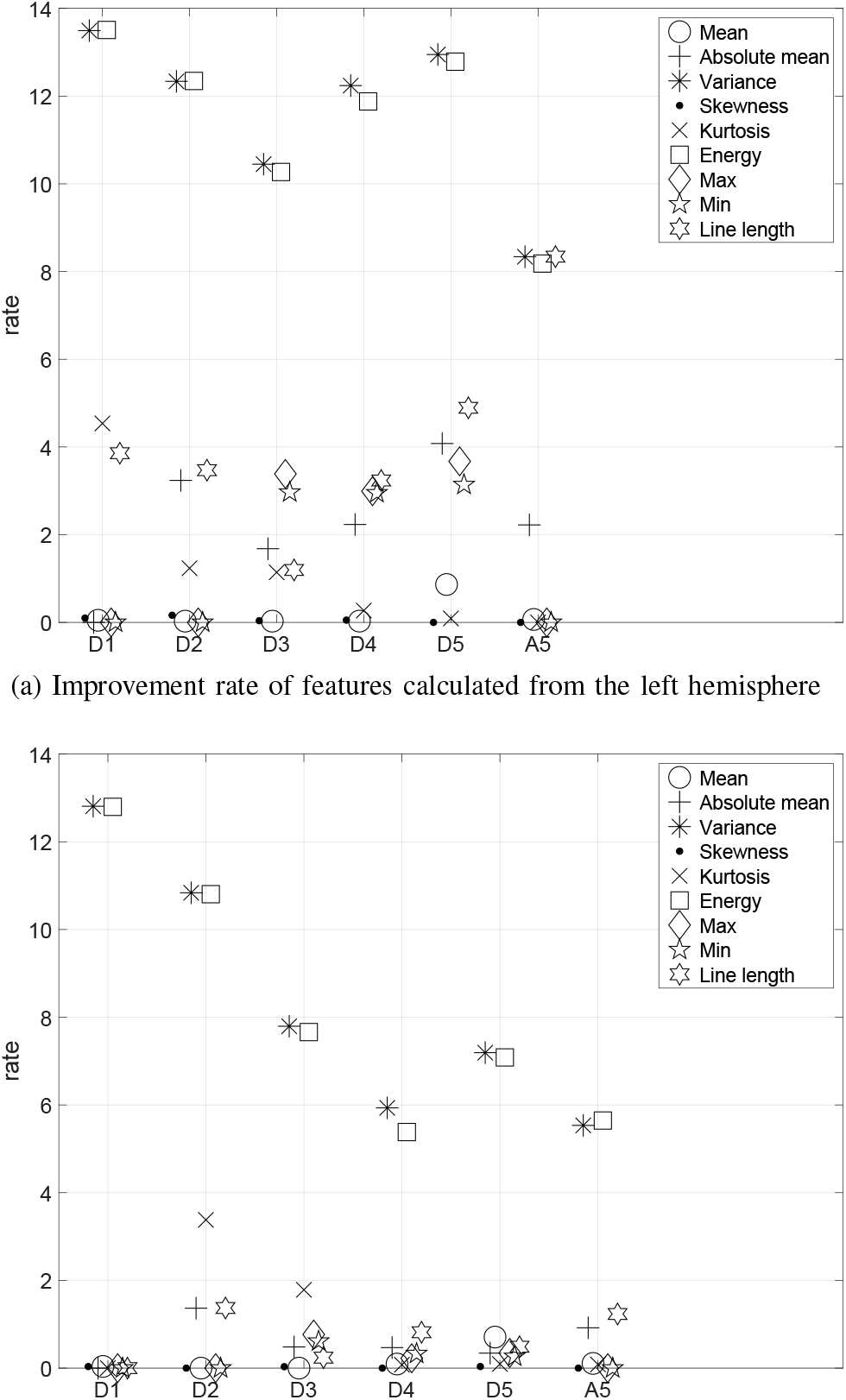
Improvement rate based on the Bayesian method of time-frequency domain features calculated from *discrete wavelet transform* using Daubechies 4 wavelet. The features are averaged over all channels from the left and right sides of the brain.

Therefore, the total number of features of each sample is 10 × 18 = 180. All dominant features are then normalized by the *z*-score normalization. Hence, the ANN model contains 1, 889 trainable parameters whereas the CNN model has 7, 945 parameters to train.

We apply 10-fold cross validation to test the proposed scheme. The following four metrics from each test fold are averaged and reported to assess the model performance:

- *Accuracy* is the number of correctly classified samples divided by the total number of sample.
- *Sensitivity* is the proportion of actual seizure samples that are detected as a seizure.
- *Specificity* measures the ratio of a number of correctly classified normal samples to the normal sample size.
- *F1-score* is a harmonic mean of positive predictive value (PPV) and sensitivity where PPV is the proportion of detected seizure samples correctly classified [32].

Figure 5 illustrates the averaged accuracy and F1-score evaluated on each EEG record. As a result, accuracies from these models are similar and are typically above 90% in every case. Apparently, CNN obtains slightly higher accuracy in many cases. They also provide similarly high specificities as shown in Figure 6. However, sensitivities and F1-score of classification of the CNN model are mostly less than of ANN. Table IV shows the averaged metrics presented in percentage over all records from the test samples. We can see that in overall CNN yielded 99.07% accuracy and 99.63% specificity whereas 98.62% accuracy and 98.92% specificity were obtained by ANN. On the other hand, ANN gave 77.04% F1-score and 82.05% sensitivity, and CNN provided 65.69% F1-score and 66.76% sensitivity.

**Fig. 5:**
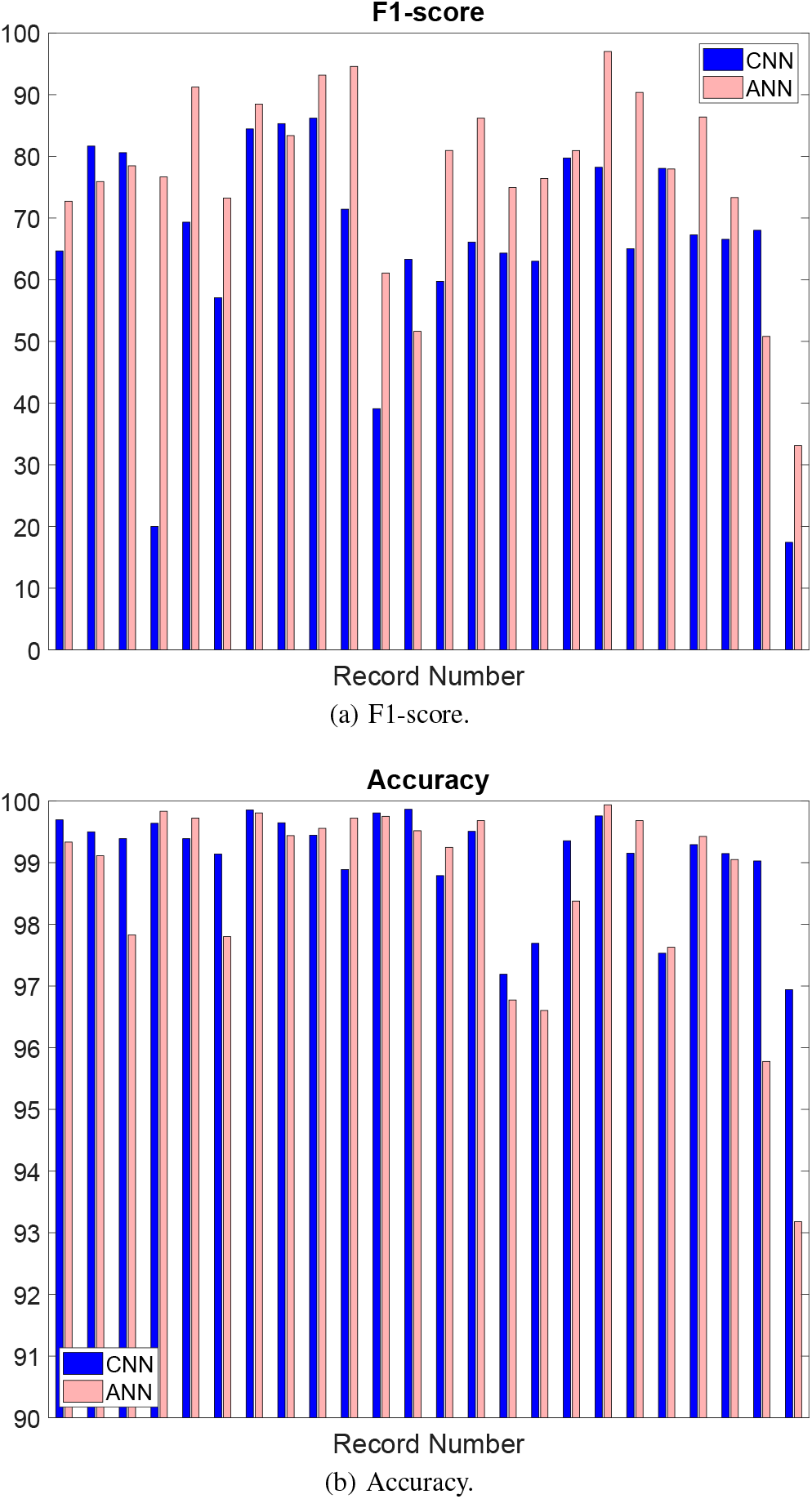
Classification performance of CNN and ANN.

**Fig. 6:**
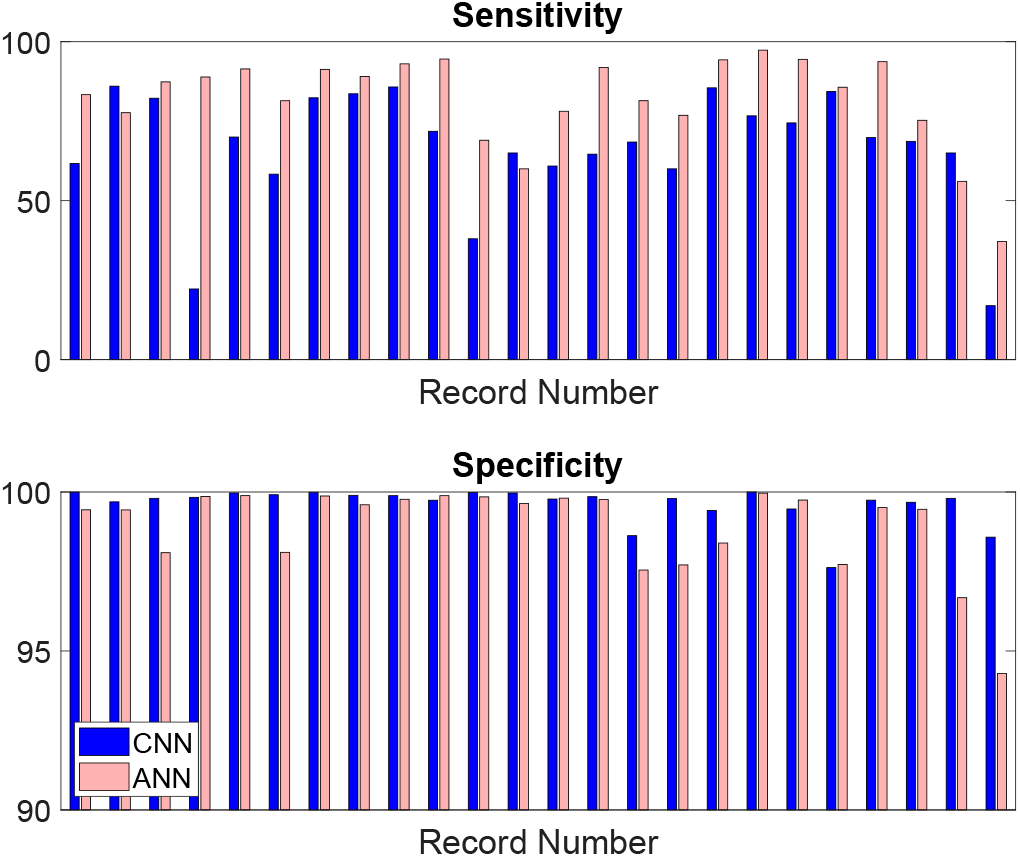
Sensitivity and specificity of classification using CNN and ANN models.

According to 15% and 11% differences of specificity and F1-score, ANN seems to be much more useful than CNN in the seizure detection that is known to be involved with imbalanced data. From the data description, there are much less seizure samples than the normal, especially some records have a small seizure duration with less than 20 seconds. It means that a few wrong detections of seizure samples can cause considerable reduction in sensitivity. For example, detecting 4 epileptic samples as normal from 40 seizure samples causes 10% difference in sensitivity. On the other hand, incorrect detections of normal samples slightly affects specificity. Therefore, 15% difference in the sensitivity and 11% in F1-score between ANN and CNN provide moderately different results in detection. This means ANN seems to detect abnormal samples better than CNN in a certain level while CNN has more potential to classify normal people. If model complexity is taken into account, these two models are comparable when achieving the same level of accuracy. Our ANN model has less parameter but requires a feature extraction process, while the CNN model has more parameters but takes multi-channel raw EEG signals as input.

**TABLE IV:** Averaged performance metrics over records and number of model parameter.

| Model | Sensitivity | Specificity | Accuracy | F1 | Parameters |
| --- | --- | --- | --- | --- | --- |
| CNN | 66.76 | 99.63 | 99.07 | 65.69 | 7,945 |
| ANN | 82.05 | 98.92 | 98.62 | 77.04 | 1,889 |

## VI. CONCLUSIONS

This research proposes a deep CNN model for classifying seizure events in multi-channel EEG signals. The model input is a raw scalp signal and the model is adjusted to reduce the complexity while achieving a promising accuracy. To compare with a deep artificial neural network (ANN), selecting significant features based on the improvement rate of Bayesian classification error is applied, and all combined prominent features are taken as the input of ANN. Performing classifications on 24 EEG recordings from CHB-MIT database shows that the accuracies of CNN and ANN are comparable (99.07% and 98.62% respectively). ANN achieves 11% higher in F1-score and has less number of parameters. However, ANN also requires 10 features calculated based on both time and frequency domains and from all EEG channels. For this reason, if we focus on the seizure detection in clinic, the ANN model with dominant features is recommended. However, if the model accuracy is primarily considered, CNN can be a favorable approach as it requires no background knowledge about ictal patterns while it can achieve a comparable overall accuracy to ANN.

## ACKNOWLEDGMENT

This research is financially supported by the 100th Anniversary Chulalongkorn University Fund for Doctoral Scholarship and the 90th Anniversary of Chulalongkorn University Fund (Ratchadaphiseksomphot Endowment Fund).

